# Phosgene Inhalation Causes Hemolysis and Acute Lung Injury

**DOI:** 10.1101/521724

**Authors:** Saurabh Aggarwal, Tamas Jilling, Stephen Doran, Israr Ahmad, Jeannette E. Eagen, Stephen Gu, Mark Gillespie, Carolyn J. Albert, David Ford, Joo-Yeun Oh, Rakesh P. Patel, Sadis Matalon

## Abstract

Phosgene (Carbonyl Chloride, COCl_2_) remains an important chemical intermediate in many industrial processes such as combustion of chlorinated hydrocarbons and synthesis of solvents (degreasers, cleaners). It is a sweet smelling gas, and therefore does not prompt escape by the victim upon exposure. Supplemental oxygen and ventilation are the only available management strategies. This study was aimed to delineate the pathogenesis and identify novel biomarkers of acute lung injury post exposure to COCl_2_ gas. Adult male and female C57BL/6 mice (20-25 g), exposed to COCl_2_ gas (10 or 20ppm) for 10 minutes in environmental chambers, had a dose dependent reduction in P_a_O_2_ and an increase in P_a_CO_2_, 1 day post exposure. However, mortality increased only in mice exposed to 20ppm of COCl_2_ for 10 minutes. Correspondingly, these mice (20ppm) also had severe acute lung injury as indicated by an increase in lung wet to dry weight ratio, extravasation of plasma proteins and neutrophils into the bronchoalveolar lavage fluid, and an increase in total lung resistance. The increase in acute lung injury parameters in COCl_2_ (20ppm, 10min) exposed mice correlated with simultaneous increase in oxidation of red blood cells (RBC) membrane, RBC fragility, and plasma levels of cell-free heme. In addition, these mice had decreased plasmalogen (plasmenylethanolamine) and elevated levels of their breakdown product, polyunsaturated lysophosphatidylethanolamine, in the circulation suggesting damage to cellular plasma membranes. This study highlights the importance of free heme in the pathogenesis of COCl2 lung injury and identifies plasma membrane breakdown product as potential biomarkers of COCl_2_ toxicity.

## Introdution

Phosgene (COCl_2_) was first synthesized in 1812 by exposing a mixture of carbon monoxide and chlorine to sunlight. It reacts slowly with water to form carbon dioxide (CO_2_) and hydrochloric acid (HCl). Phosgene is a widely used in industry for the synthesis of isocyanate-based polymers, carbonic acid esters and acid chlorides. It is also used in the manufacture of dyestuffs, some insecticides and pharmaceuticals and in metallurgy. Although most COCl_2_ is produced on-site, a fair amount is transported by rail or truck making transportation accidents likely (19; 25; 37).

Like the halogens chlorine (Cl_2_) and bromine (Br_2_), COCl_2_ is cheap and produced easily; however, unlike Cl_2_ and Br_2_ it cannot be detected by smell in small but dangerous concentrations, increasing the probability that people may be exposed to potentially harmful concentrations before they can react (19). As an example, a severe accident involving release of COCl_2_ occurred in a DuPont Factory in Belle, West Virginia in 2010 resulting in worker fatalities https://www.csb.gov/dupont-corporation-toxic-chemical-releases/. Phosgene may also be generated during welding and during fires involving plastics and solvents containing Cl_2_ placing workers and first responders at serious risk (53). Phosgene exposure resulted in more than 80% of all chemical-related deaths in WWI (25) and it may have been used against civilian targets as recently as 2017 and 2018 in Syria http://www.worldinwar.eu/the-gas-used-in-khan-shaykhun-probably-the-phosgene-syria/.

Finally, chloroform (CHCL_3_), a former anesthetic agent, is metabolized to COCl2 in the liver and the kidneys (7; 42), which may account for some of the well-known CHCL_3_ toxicity Presently, COCl_2_ injury to the mammalian blood gas barrier and systemic organs has not been rigorously documented. Cl_2_ and Br_2_, which mainly interact with small molecular weight antioxidants present in the epithelial lining fluid as well as lipids and proteins on the surface of airway and epithelial cells (51; 56). We have also demonstrated that Cl_2_ and Br_2_ react with plasmalogens, present in all cell membranes and the epithelial lining fluid, as well as glutathione, to form halogenated adducts (fatty acids and aldehydes), which were detected in the alveolar space, lung tissues and plasma (10; 15) and may be responsible for the mediation of injury to distal organs, such as the heart (3). Whether COCl_2_ reacts with plasmalogens is not known but there is evidence regarding its reaction with phospholipids **i**n vivo in CHCl_3_ hepatotoxicity as well as with phosphatidylcholine in vitro (12; 18). In addition, because of its slow hydrolysis rate and solubility, COCl_2_ may cross the blood-gas barrier, enter the capillary circulation and damage RBCs directly (44; 48). Interestingly, humans exposed to COCl_2_ show transient declines in RBC count (40). If this decline is due to hemolysis, free heme/hemoglobin mediated processes may underlie tissue injury after COCl_2_ exposure. Indeed, one study showed that exposure of rats to COCl_2_ increased hemolysis (48).

Thus the purpose of this study was to vigorously assess whether COCl_2_ induced injury to the blood gas barrier and red blood cells of unanesthetized mice. To accomplish these goals, we exposed equal numbers of adult male and female mice to 10 or 20 ppm COCl_2_ in environmental chambers for 10 min and returned them to room air, and assessed the following parameters at different times post-exposure: survival, body weight changes, physiological and histochemical indices of lung function and injury including arterial blood gases, plasmalogen levels, and levels of hemoglobin and free heme in the plasma. We show that mice exposed to COCl_2_ showed evidence for increased RBC fragility, oxidative injury to RBC membranes, damage to plasma plasmalogens and the onset of delayed but severe lung injury which mimics human Adult Respiratory Distress Syndrome (55). Furthermore, there was considerable oxidation of important RBC structural proteins and band 3 which may contribute to hemolysis. The results of these studies pave the road for additional studies for the development of countermeasures against hemoglobin and heme, such as haptoglobin and hemopexin, as novel countermeasures for COCl_2_ injury.

## Materials and Methods

### Reagents

Ketamine was obtained from Vedco Inc. (St. Joseph MO); Xylazine from Vet One, (Boise, ID); the heme assay kit was from QuantiChrom (Product No. DIHM-250; BioAssay Systems, Hayward, CA); 4mm Pyrex solid glass beads from Sigma-Aldrich (St Louis, MS); 4-20% Tris·HCl Criterion precast gels from Bio-Rad Laboratories (Product number: 567-1094, Hercules, CA); Amido Black from Sigma-Aldrich (St Louis, MS); RIPA buffer from Thermo Fisher Scientific (MA); Oxyblot protein oxidation detection kit from EMD Millipore (Product number: S7150, Billerica, MA)

### Animals

Adult male and female C57BL/6 mice (20-25g) were bought from Charles River (Wilmington, MA). Mice were allowed to acclimatize in the Animal Vivarium located in the basement of the Biomedical Research Building II for at least four days, where they were cared for by personnel from Animal Resources Program. All animal care and experimental procedures were approved by the Institutional Animal Care and Use Committee at the University of Alabama in Birmingham (protocol # 21451). Mice that exhibited respiratory symptoms, or refused to eat and drink, were not included in these studies.

### Exposure to Phosgene Gas

On the day of the exposure, mice of either sex were placed, five at a time, in a 4.5 L glass exposure chamber ((1; 2)). The exposure chambers were located inside a negative pressure hood, inside a room at the Animal Vivarium, maintained at negative pressure compared to the rest of the Vivarium

The exposure chamber was connected to either compressed air or COCl_2_ gas (nominal concentrations of 20 or 10 ppm, certified within 2%, purchased from SpecGas, Inc. Warminster, PA). The flow rate was set at 5 L/min. In some cases, the concentration of COCl_2_ was monitored by an Analytical Technology, Inc. (Collegeville, PA) A21 Gas Sampling System, F12D Gas Transmitter with Phosgene Sensor (00-1016) Standard Range = 0-100 ppm; Resolution = 0.1 ppm). In addition, the concentration of CO_2_ in the chambers was monitored by a UEi C20 combustion meter (Beaverton OR). The gas exiting the chamber was passed through a 10% solution of NaOH to scavenge the COCl_2_ and vented to the roof of the building.

At the end of each ten min exposure, the gas was turned off, the chamber lid was removed and after a short period of time, the mice were returned to their cages in the Vivarium where they were provided with food and water *ad* libidum and observed by both laboratory and technicians of the Animal Facility. All personnel involved in the exposure of mice to COCl_2_ underwent special training by the Occupational and Safety division of the University of Alabama at Birmingham. A COCl_2_ detector, (purchased from GasSensing, Hull IA), equipped with an audible alarm and a strobe light was mounted on the wall of the exposure room and was set to sound an alarm if the COCl_2_ concentration in the room exceed 0.5 ppm COCl_2_.

### Histological analysis of the mouse lung

Mice were euthanized with an intra-peritoneal injection of ketamine and xylazine (160 and 80 mg/kg body weight, respectively). The chest was opened and the lungs were removed, fixed in 10% formalin for 24 hours and dehydrated in 70% ethanol before embedding in paraffin. Paraffin-embedded tissues were cut into 4 μm sections then de-paraffinized and rehydrated using Citrisolv (d-limonene based solvent) and isopropanol, respectively. The sections were stained with hematoxylin and eosin (H & E). Images were captured using a Leica DMI 6000 B microscope (Leica Microsystems Inc., Bannockburn, IL) and Leica Application Suite V4.2 software.

### Plasma heme assays

Heme levels were measured in plasma samples by two different methods: first, by using the QuantiChrom heme assay kit (Product No. DIHM-250; BioAssay Systems, Hayward, CA), according to the manufacturer’s instructions; and second, by spectral deconvolution with least square fitting analyses (45)

### Red Blood Cell Fragility

Red Blood Cells from air and COCl_2_ exposed mice were washed thoroughly to remove free heme. RBCs were re-suspended, at 1.0% hematocrit, with 4mm Pyrex solid glass beads (10 beads, 0.4 ml RBC suspension volume in 2 ml round bottom Eppendorf tubes) in DPBS. This solution was rotated 360° for 2 hours at 24 rpm at 37°C. The hemoglobin released from the RBCs during rotation was transferred into a new tube and centrifuged at 13,400g for 4 min and the absorbance of the supernatant were recorded at 540nm.

### Respiratory mechanics

Mice were mechanically ventilated and challenged with increasing concentrations of methacholine as described previously (59). Briefly, mice were anesthetized with diazepam (17.5 mg/kg) and ketamine (450 mg/kg), intubated, connected to a ventilator (FlexiVent; SCIREQ, Montreal, PQ, Canada) and ventilated at a rate of 160 breaths per minute at a tidal volume of 0.2 ml with a positive end-expiratory pressure of 3 cm H_2_O. Total respiratory system resistance (R) and elastance (E) were recorded continuously as previously described (*ibid.*). Baseline lung volume was set via deep inhalation. Increasing concentrations of methacholine chloride (0–50 mg/ml, Sigma-Aldrich, St Louis, MS) were administered via aerosolization within an administration time of 10 sec. Airway responsiveness was recorded every 15 seconds for 3 minutes after each aerosol challenge. Broadband perturbation was used and impedance was analyzed via constant phase model.

### Arterial blood gases

Mice were anesthetized with Isoflurane (5% for induction, 2% for maintenance) using compressed air as vehicle. The abdomen was opened, the mesentery was externalized to the left side of the mouse to visualize the abdominal aorta. Arterial blood was collected into a heparinized syringe through a 23 gauge needle inserted into the aorta. Blood gas analysis was performed immediately after collection using an Element POC analyzer (Heska, Loveland, CO).

### SDS-PAGE and Oxyblots

Mice were anesthetized and euthanized. Their lungs were then lavaged with one ml of NaCl instilled and withdrawn three times. The cells were pelleted by centrifugation (10 min at × 200g). Cleared supernatants were used to measure the protein concentration by the BCA assay and equal volume of BALF (2μl) were loaded on 4-20% Tris·HCl Criterion precast gels (Product number: 567-1094, Bio-Rad Laboratories, Hercules, CA). In addition, RBCs were separated from the plasma and hemolyzed with 20 mM hypotonic HEPES (4-(2-hydroxyethyl)-1-piperazineethanesulfonic acid) buffer. The mixture was centrifuged at 14000xg for 20 min and the RBC pellet was dissolved in RIPA (Radioimmunoprecipitation assay) buffer. The protein was quantified by the BCA method and equal amounts of proteins (10μg) were loaded into a 4-20% gradient gel and proteins were separated and stained with Amido Black.

The presence of protein carbonyl groups was assessed using the Oxyblot protein oxidation detection kit (Product number: S7150, EMD Millipore, Billerica, MA), according to the manufacturer’s protocol (2). Briefly, the carbonyl groups in the protein side chains were derivatized to 2,4-dinitrophenylhydrazone by reacting with 2,4-dinitrophenylhydrazine. Precisely, 10 μg of protein was used for each sample, and the 2,4-dinitrophenol-derivatized protein samples were separated by polyacrylamide gel electrophoresis, as described previously. Polyvinylidene fluoride membranes were incubated for 1 hour in the stock primary antibody (1:150 in 1% PBS/TBST buffer), and after washing, for 1 hour in the stock secondary antibody (1:300 in % PBS/TBST buffer). Membranes were washed 3× in TBST and visualized, as described previously. The abundance of protein carbonylation was assessed by densitometry of each lane and normalization for each lane protein loading was done by SDS PAGE gel quantification.

### Quantification of plasmalogens in plasma

Equal number of male and female mice were sacrificed 24 h post exposure to either air or COCl_2_. Plasma was subjected to a modified Bligh-Dyer lipid extraction (6) in the presence of lipid class internal standards including 1-0-tetradecanoyl-sn-glycero-3-phosphoethanolamine and 1,2-ditetradecanoyl-sn-glycero-3-phosphoethanolamine (9). Lipid extracts were diluted in methanol/chloroform (1/1, v/v) and molecular species were quantified using electrospray ionization mass spectrometry on a triple quadrupole instrument (Themo Fisher Quantum Ultra) employing shotgun lipidomics methodologies (22). Ethanolamine glycerophospholipid and lysophosphatidylethanolamine molecular species were first converted to 9-fluorenylmethoxycarbonyl (fMOC) derivatives and then quantified in the negative ion mode using neutral loss scanning for 222.2 amu (collision energy = 30eV).

### Statistical Analysis

Statistical analysis was performed using GraphPad Prism version 4.01 for Windows (GraphPad Software, San Diego, CA). The mean ± SEM was calculated in all experiments, and statistical significance was determined by either the one-way or the two-way ANOVA. For one-way ANOVA analyses, Tukey’s post-test post-hoc testing was employed, while for two-way ANOVA analyses, Bonferroni post-tests were used. Overall survival was analyzed by the Kaplan-Meier method. Differences in survival were tested for statistical significance by the log-rank test. A value of P < 0.05 was considered significant.

## Results

### Whole body exposure to COCl_2_ in glass chamber

As shown in **Figure 1**, COCl_2_ concentrations in the exposure chamber rose after onset of COCl_2_ flow and reached a steady state after approximately five min. There were no differences in the concentration profiles whether or not mice were present in the chambers, indicating that absorption of COCl_2_ by the fur was negligible. The differences in steady state concentrations recorded by the COCl_2_ detector and the certified nominal COCl_2_ concentrations were within the stated accuracy of the detector. Based on the profiles shown in **Figure 1**, the areas under the concentration profiles were 57.5 and 156 *ppm x min*. For simplicity we are referring to the exposures as 10 and 20 ppm. CO_2_ concentrations in the exposure chamber were undetectable during the exposure period.

**Figure 1.**
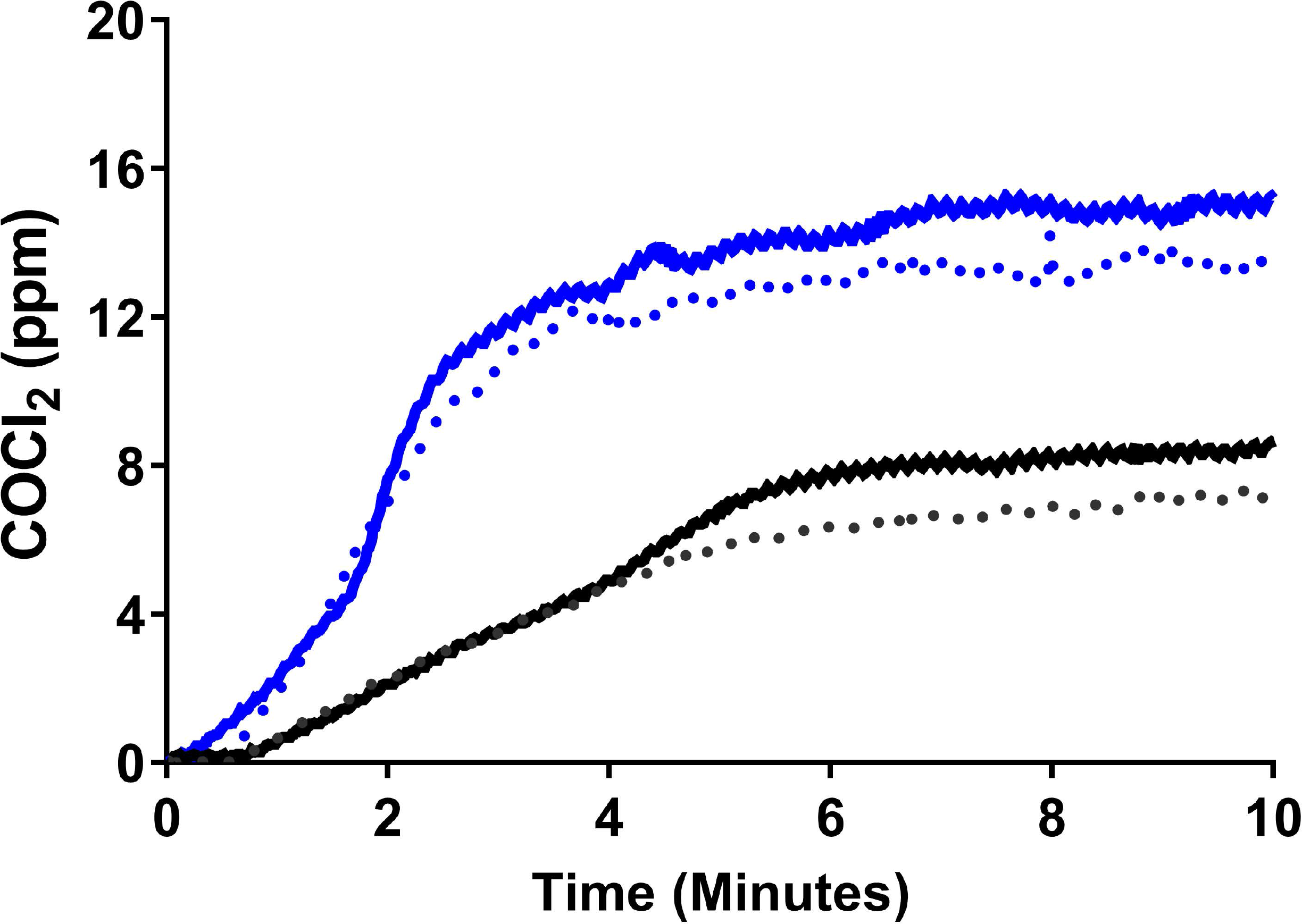
Continuous measurement of COCl_2_ concentrations inside the exposure chambers. COCl_2_ gas (nominal concentrations of 20 ppm (blue line) or 10 ppm (black lines) flowed into the exposure chambers (volume=4.5 L) at 4.4 L/min. The concentration of COCl_2_ was monitored by a. Analytical Technology, Inc. (Collegeville, PA) A21 Gas Sampling System, F12D Gas Transmitter with Phosgene Sensor. Measurements were conducted with five mice in the chamber (dotted lines) or with empty chambers (solid lines). Measurements were repeated twice with similar results.

### Exposure to COCl_2_ results in hypoxemia and respiratory acidosis

Adult C57BL/6 mice were exposed to air to COCl_2_ (10ppm or 20ppm) for 10 minutes. Mice appeared alert and unaffected by COCl_2_ both during the exposure and within the first 12 hours of return to air. After that time till they were sacrificed 24 hours later they exhibited reduced activity levels but they did not exhibit overt respiratory distress, i.e., lacked labored breathing, flaring of the nostrils and expiratory grunting. Measurements of arterial blood gases **(Figure 2)**, Pa_O2_ **(2A)**, Pa_CO2_ **(2B)**, pH **(2C)**, and 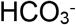 **(2D)** was performed in anesthetized mice using an Element POC analyzer (Heska, Loveland, CO) as described previously (27). Twenty four hours post COCl_2_, mice developed dose-dependent hypoxemia and respiratory acidosis, despite increases in plasma bicarbonate. The increase in 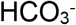 was consistent with a decrease in plasma chloride concentration (data not shown).

**Figure 2.**
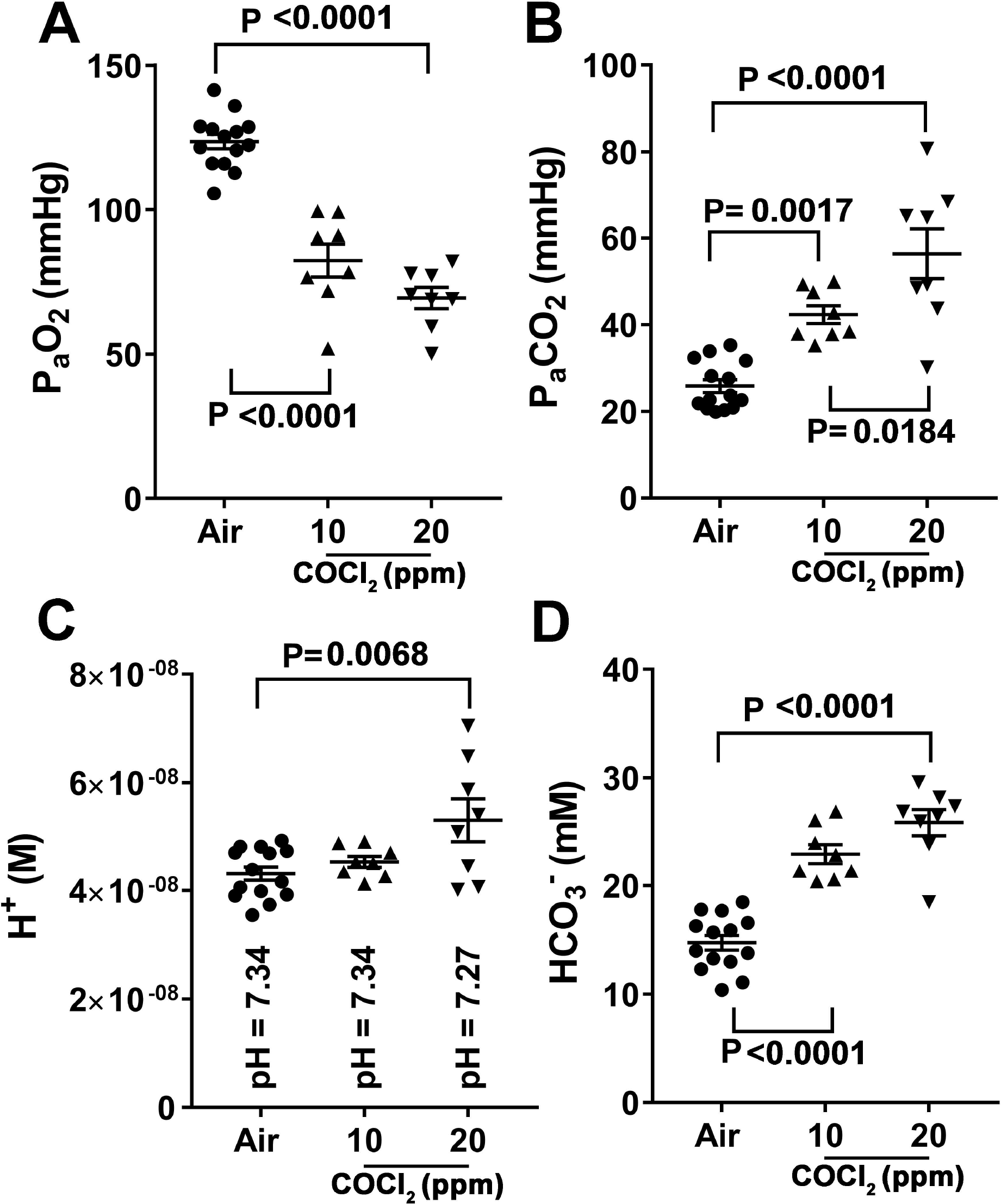
Measurements of arterial blood gases in COCl_2_ exposed mice. Equal number of male and female mice were anesthetized at 24 h post exposure with isoflurane, and one ml blood sample was withdrawn from the aorta, following opening of the chest cavity. PO_2_, PCO_2_, pH and 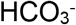 were measured with an Element POC analyzer (Heska, Loveland, CO). Values are means ± 1 SEM. Each point represents a different mouse. Statistical analysis was performed by one way analysis of variance followed by Tukey’s post-hoc testing.

### Exposure to COCl_2_ increases mortality in mice

As shown in **Figure 3**, air breathing mice continued to gain weight, eat and drink for the next 21 days **(3A)**. Mice exposed to 10 ppm COCl_2_ experience a transient 15% decrease of body weight during the first 24 h post exposure, but resumed eating and drinking and gain weight as the air controls **(3B)**. All of these mice were alive at 21 d post exposure. On the other hand, mice exposed to 20 ppm COCl_2_ for 10 min and returned to room air, continued to lose weight during the first 5 days post exposure **(3C)**. Two of these mice died during the first 24 h post exposure and three mice were sacrificed because their body weights dropped below 30% of their pre-exposure weights. Kaplan-Meyer curves for mice exposed to 10 or 20 ppm COCl_2_ are shown in **Figure 3D**. Mice exposed to 20 ppm for 10 min COCl_2_ exhibited approximately 50% survival, with all deaths occurring in the first 6 days post exposure.

**Figure 3.**
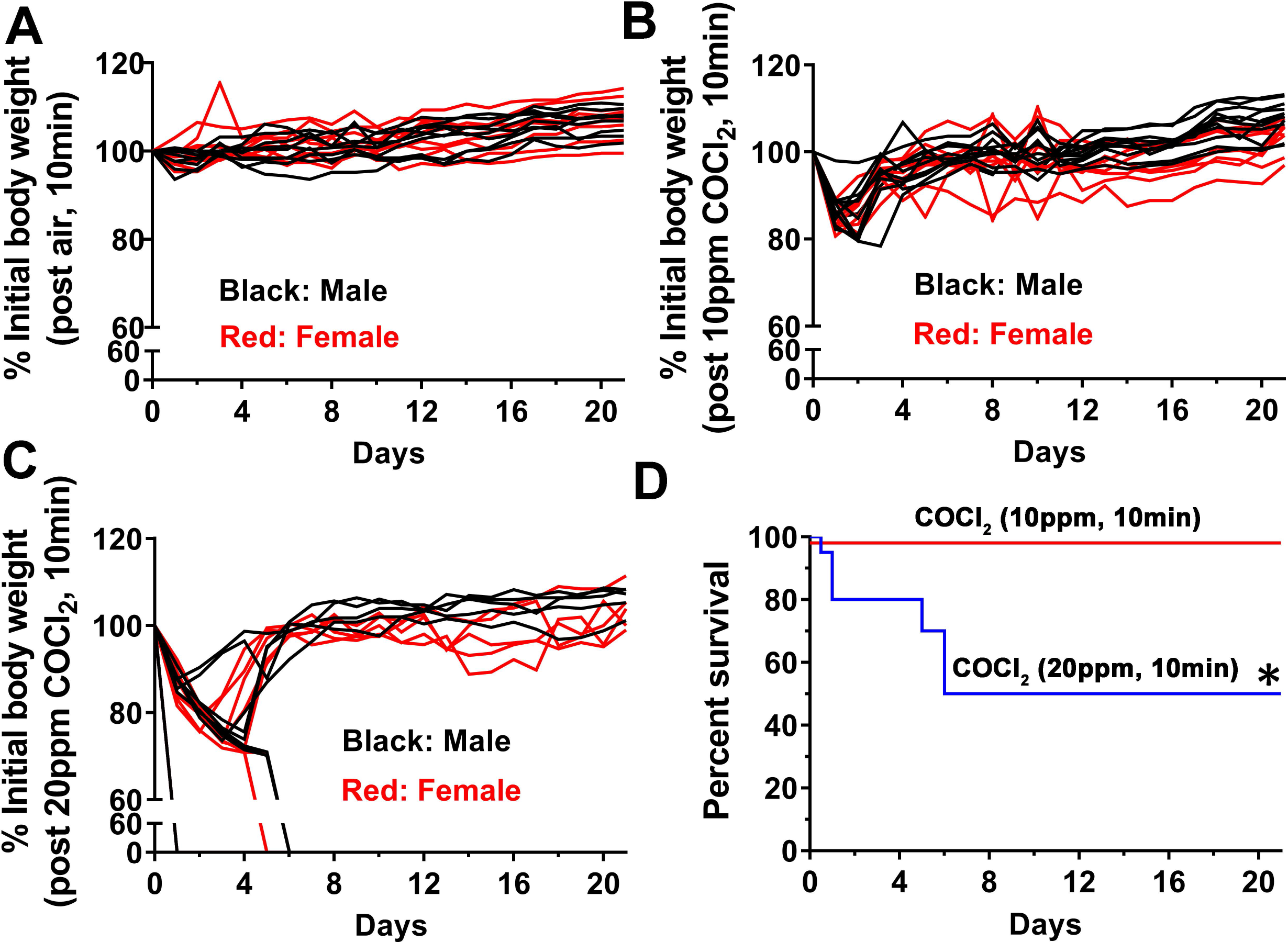
Measurements of body weight and mortality in mice exposed to COCl_2_. Equal number of male and female C57BL/6 were placed five at a time in the environmental exposure chambers and exposed to air (n=18), 10 ppm COCl_2_ (n=20) or 20 ppm COCl_2_ (n=20) for 10 min. At the end of the exposure, the gas flow was stopped, the top lid of the chamber removed and the mice were transferred to their normal holding cages. Their body weights were measured daily and expressed as % of initial weights. Mice exposed to air (**A**) continued to gain weight; mice exposed to 10 ppm COCL_2_ for 10 min (**B**) lost no more than 20% of their initial body weight in the first 24 hours but gained weight thereafter. Mice exposed to 20 ppm for 10 min lost a significant higher levels of their body weights (**C**). Two male and female mice each were found dead in their cages one day post exposure. The sharp drops of body weight shown in **3C** indicate that the mice were either sacrificed due to a loss of more than 30% of body weight or were found dead. No significant difference among male and female mice was seen. Equal numbers of male and female mice were exposed in each case (n=10). Kaplan-Meyer survival curves (**D**) for 10 ppm COCl_2_ (n=20; red line) or 20 ppm COCl_2_ (n=20; blue line) for 10 min (equal numbers of male and female mice) *= p=0.0006 as compared to air or 10 ppm COCl_2_ according to the Long-rank (Mantel-Cox) test. All mice exposed to air were alive at 21 days post exposure

### Exposure to 20 ppm COCl_2_ damages the distal lung regions and results in inflammation

Consistent with the data shown in **Figure 3**, mice exposed to COCl_2_ (20 ppm, 10 min) had severe injury to the blood gas barrier 24 hours post exposure **(Figure 4)**. There was a progressive and significant increase of the lung/wet dry weight **(4A)**, consistent with the presence of interstitial and alveolar edema, and a 10 fold increase of plasma proteins in the bronchoalveolar lavage fluid (BALF) **(4B)**. Interestingly, the magnitude of increased lung edema far exceeded (by 5-10fold) that observed in mice exposed to HCl (59), Cl_2_ or Br_2_ (1; 2; 29); maximal BALF protein levels in these are in the 1-2mg/ml range. SDS PAGE analysis of cell free BALF showed a large increase in a number of protein bands, including a 67 kDA band, which is the expected size for albumin **(4C)**. It should be noted that the pattern observed in SDS-PAGE gels was similar to that of plasma (far right two lanes in **Figure 4C**), indicating complete breakdown of the alveolar barrier.

**Figure 4.**
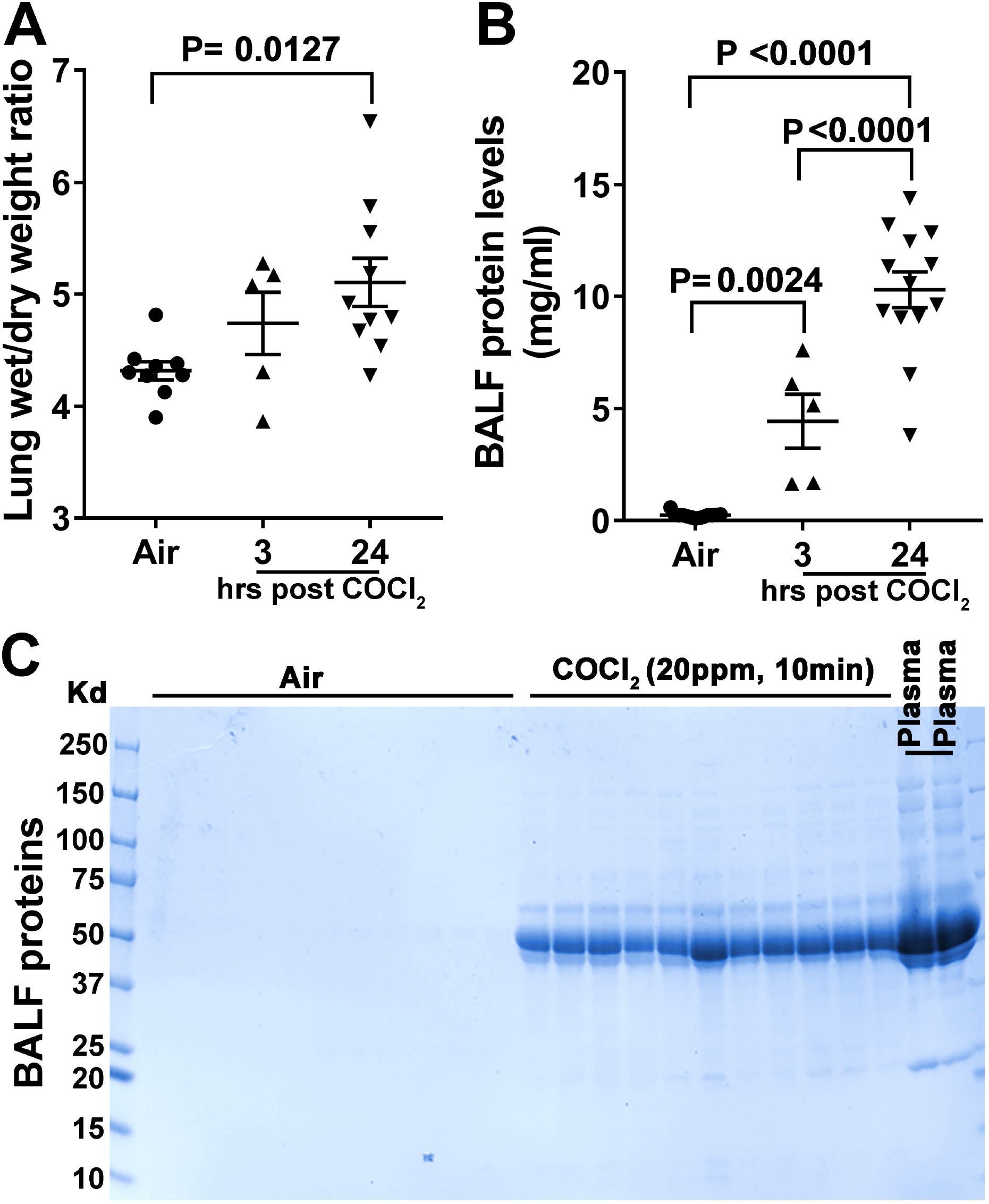
Measurement of lung edema in mice exposed to COCl_2_. Equal number of male and female mice were sacrificed 3 or 24 hours post exposure to either air or COCl_2_; their chests were opened and lungs removed *en block*, blotted and non-lung tissue was removed. They were then weighted for determination of wet weight and placed in an oven at 70 °C for seven days at which point they were reweighted for measurement of dry weights. Data shown are wet/dry weights **(A)**. Mean±1 SEM; n=9 air; n=5 for 3 hours post COCl_2_, and n=10 for 24 hours post COCl_2_; In a separate set of animals mice lungs were lavaged with 1 ml of NaCl. The cells were pelleted by centrifugation (10 min at ×200g) and protein in the cell free lavage was determined by the BCA method **(B)**. Mean±1 SEM; n=9 air; n=5 for 3 hours post COCl_2_, and n=10 for 24 hours post COCl_2_. BALF proteins were also separated by SDS-PAGE (2 μl BAL). No significant amount of proteins were detected in the BALF of air breathing mice; however, significant amounts of a protein with a molecular weight of around 70 kDa (most likely albumin) was detected in the BALF of mice harvested 24 hours post COCl_2_ exposure **(C)**. Large number of other plasma proteins were also seen. Statistical analysis was done by ANOVA with Tukey’s post-test (**A**, **B**).

The total cell count remained unchanged at both 3 and 24 h post exposure (data not shown). However at 24 hours post exposure **(Figure 5)** there was a significant increase of neutrophils **(5A)** and a concomitant decrease of alveolar macrophages **(5B)**. H&E staining of the lungs of mice sacrificed at 24 hours post COCl_2_ (20 ppm for 10 min) showed that most alveoli were filled with exudate exhibiting eosin staining, indicative of high protein content **(5C),** consistent with data shown in **Figure 4**. In contrast, H&E stains of the distal lung regions of mice exposed to air, were free of exudate and showed normal architecture. Macroscopic examination of the lungs of COCl_2_ exposed mice showed severe hemorrhage **(5C)**.

**Figure 5.**
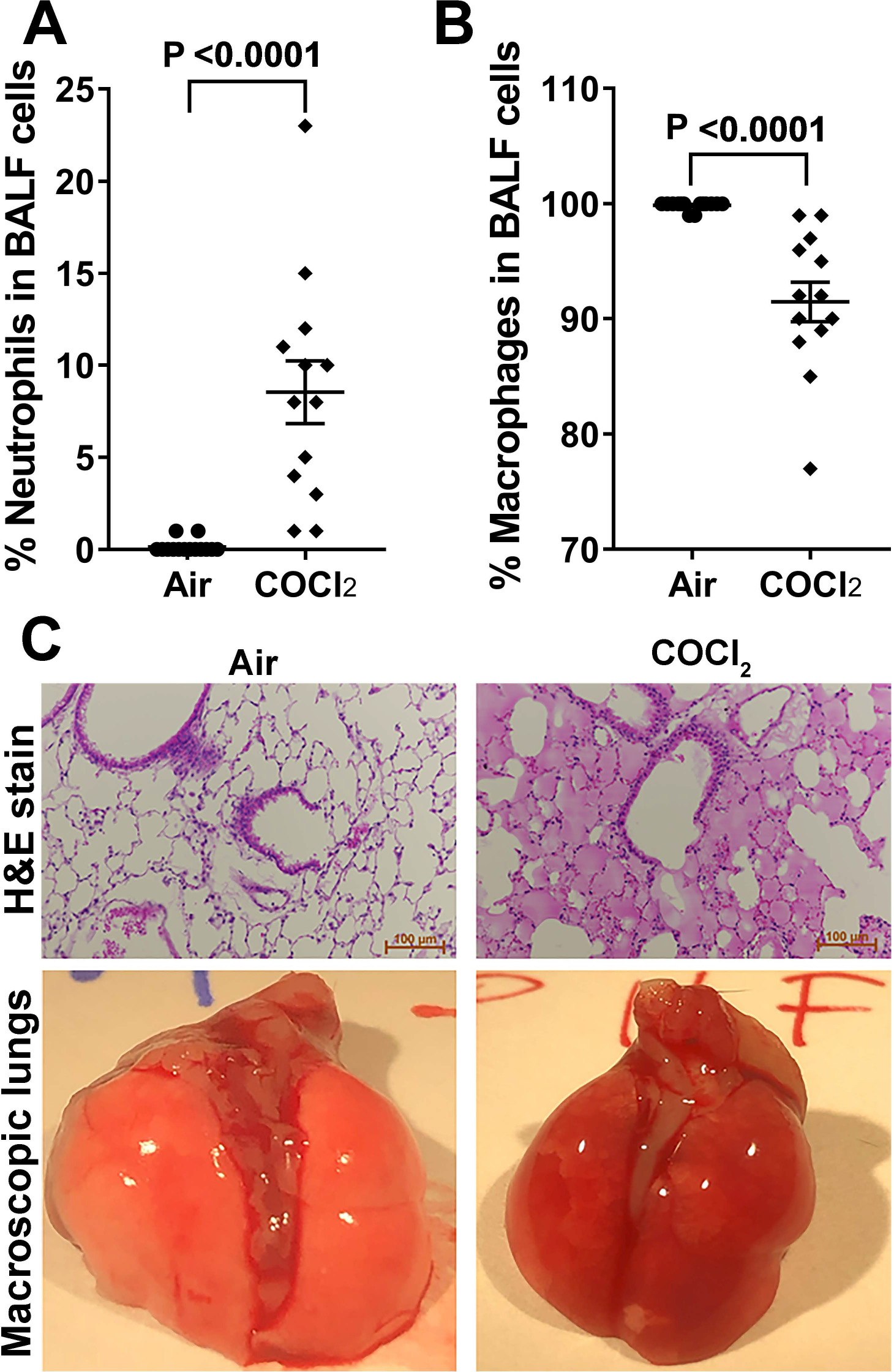
Measurements of distal lung injury in mice exposed to COCl_2_. Equal number of male and female mice were sacrificed 24 h post exposure to either air or COCl_2_. Neutrophils **(A)** and macrophages **(B)**, identified by their characteristic shapes were counted in the BALF with a hemocytometer and expressed as % of total cells. Means±1 SEM; n=14 air; n=13 COCl_2_; statistical analysis by unpaired t-test. In addition, the lungs were fixed with formalin at a 25 cmH_2_O constant pressure. Thin sections were cut from all lobes, fixed and stained with hematoxylin and eosin **(C)**. The characteristic pictures show that the lungs in COCl_2_ mice were filled with fluid containing proteins and cells. A small number of alveoli appear empty of fluid and were responsible for gas exchange. In addition, macroscopic examination of the lungs showed significant hemorrhage in COCl_2_ exposed mice **(C)**.

### Exposure to 20 ppm COCl_2_ increases airway resistance

Measurements of airway resistance and elastance with flexiVent showed a large increase of baseline airway resistance and a concomitant decrease of dynamic elastance at 24 hours post exposure (**Figure 6**). Following methacholine challenge, airway resistance **(6A)** and elastance **(6B)** changed similarly in both COCl_2_ and air exposed mice; these data indicate that in spite of the large increase of baseline airway resistance, airways did not become hyperreactive following exposure to COCl_2_, in contrast to what seen in Cl_2_ and Br_2_ exposed mice. Surprisingly, Newtonian airway resistance, an index of injury to upper airways, did not change in the COCl_2_ group indicating that the increase of airway resistance was mainly due to damage to small airways.

**Figure 6.**
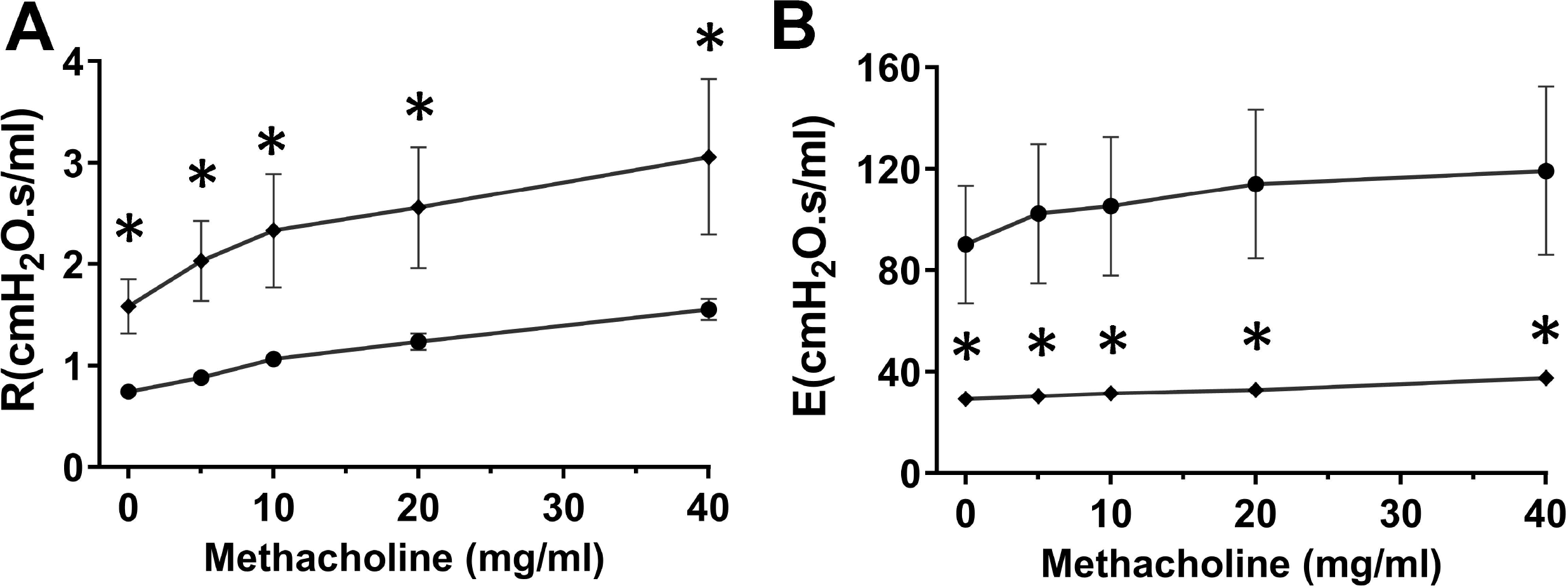
Measurements of airway resistance and elastance in mice exposed to COCl_2_. Male only C57BL/6 mice were exposed to either air (n=10; black circles) or COCl_2_ (20 ppm for 10 min; n=5; black diamonds) and returned to room air for 24 hours. They were then anesthetized and connected to a flexiVent for measurements of Airway Resistance **(A)** or Elastance **(B)**. Numbers are means ± 1 SEM; *p<0.05 (unpaired t-test). * p<0.05 by Student’s t-test.

### Exposure to COCl_2_ damages red blood cells and causes hemolysis

First, RBC membranes were analyzed for the presence of oxidative damage **(Figure 7)**. For these studies, RBC were collected and membranes prepared as outlined in methods. The measurement of membrane associated carbonyl adducts **(7A, B)**from RBC collected 24 hours post exposure to COCl_2_ (20 ppm, 10min) showed significantly higher levels of oxidation **(7A, B)**of a 90-95 kDa protein as well as a number of higher molecular weight proteins. Due to the limited repertoire of proteins in RBC membrane, the 90-95 kDa protein is likely to be band 3 (21); notice that both 4.1a and 4.1b bands are clearly visible and their ratio is about 1:1 in most of the preparation, as previously reported (17); the high molecular species are likely to be ankyrin and spectrin. SDS-PAGE of RBC ghosts revealed no difference in total levels of these proteins (data not shown).

**Figure 7.**
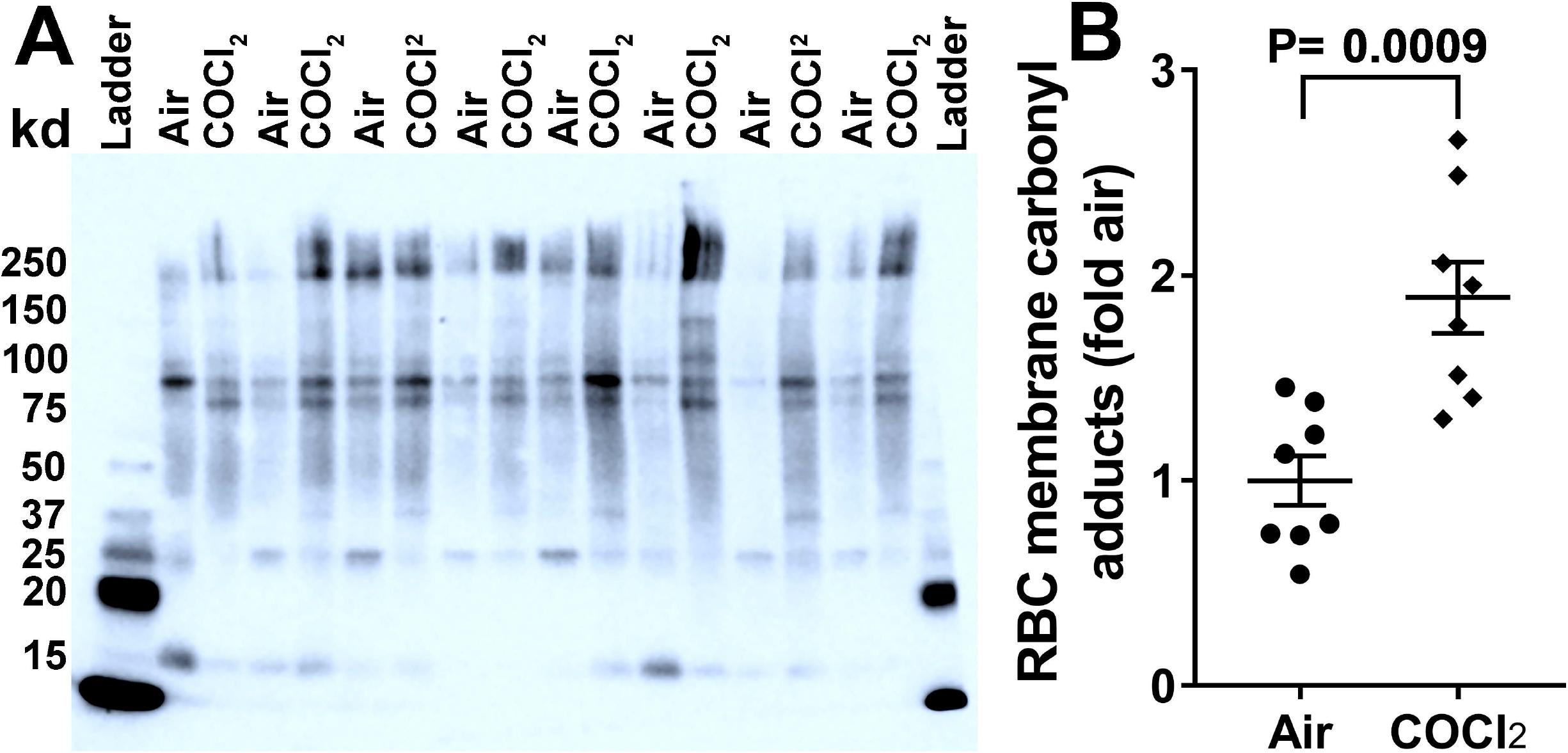
Carbonyl adducts are increased in RBC ghosts post COCl_2_ exposure. Mice were exposed to air or COCl_2_ (20 ppm, 10min). RBCs were separated from the plasma and hemolyzed with 20 mM hypotonic HEPES Buffer 24 hours later. The mixture was centrifuged at 14000 xg for 20 min and the resulting RBC pellet was dissolved in RIPA buffer. The protein was quantified by the BCA method and the presence of protein carbonyl groups was assessed using the OxyBlot Protein Oxidation Detection Kit according to the manufacturer’s protocol. Significant oxidation was noted in a protein around 95 KDa. Oxidation of high molecular weight proteins (around 250 kDa; presumably ankyrin and spectrin) was also observed **(A)**. Each lane represents a different mouse. Additionally equal amounts of proteins were loaded into a 4-20% gradient gel and proteins were separated and stained with Amido Black (data not shown). There was no difference in protein levels between the air and the COCl_2_ exposed mice. Panel **B** shows the quantification of carbonyl adducts. Values are means ± 1 SEM; Student’s t-test.

Next, the analysis of RBC fragility, showed that exposure to mechanical stress (rotation of RBCs with glass beads for 2 hours) caused significantly more hemolysis in the RBCs obtained from the mice exposed to COCl_2_ (20 ppm for 10 min) than the air exposed mice **(Figure 8A)**. Next, we measured total levels of non-encapsulated hemoglobin and heme, using an ELISA method which does not discriminate between the two. There was almost a doubling of non-encapsulated heme at 24 hours post 20 ppm COCl_2_ for 10 min **(8B)**. Measurements of total non-encapsulated heme using a spectrophometric technique followed by spectra deconvolution revealed similar increase in total non-encapsulated heme **(8C)** and a significant increase if free heme, formed by the oxidation of hemoglobin **(8D)**.

**Figure 8.**
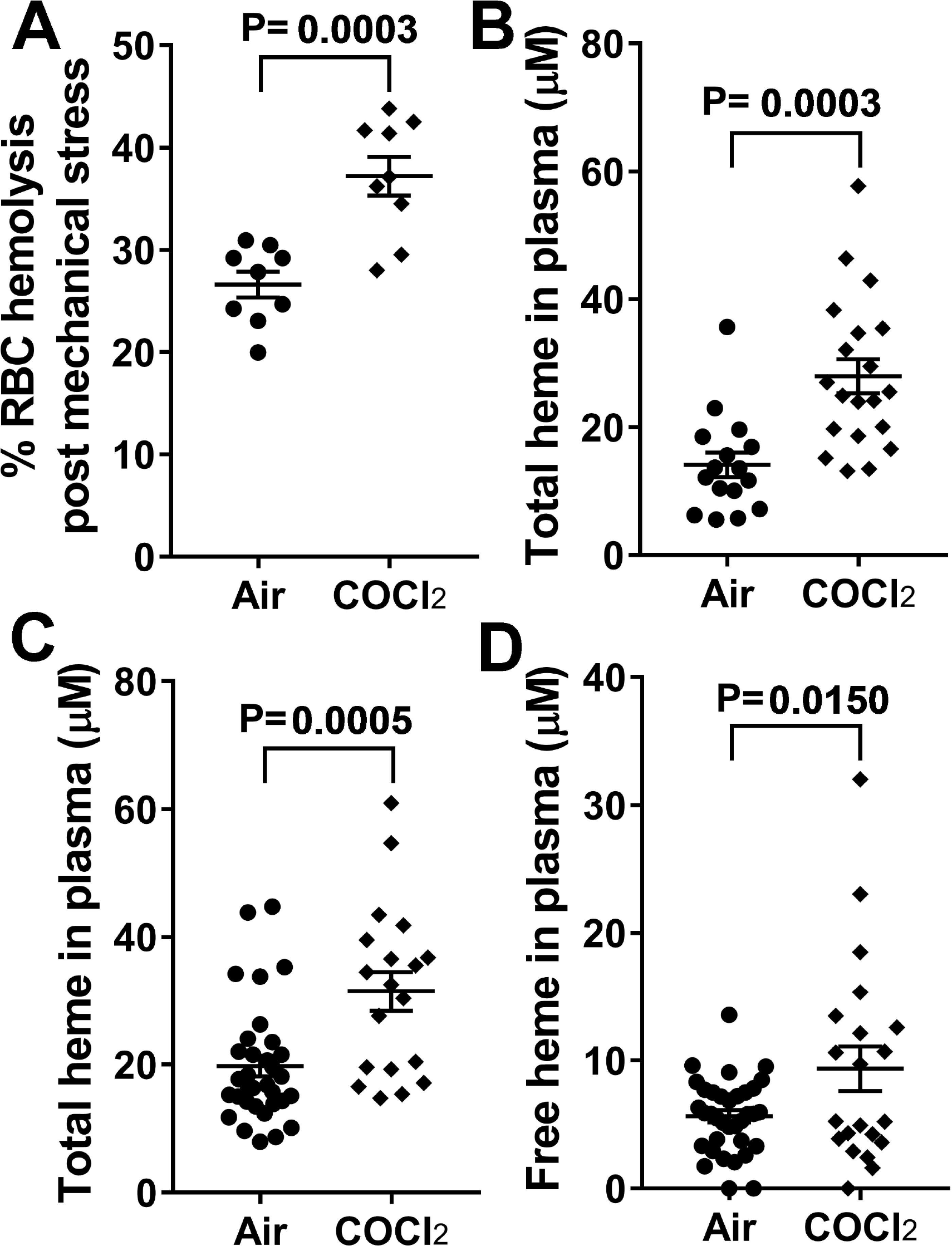
COCl_2_ exposure increases RBC fragility in mice. Equal number of male and female mice were sacrificed 24 hours post exposure to either air or COCl_2_ (20 ppm for 10 min). Blood was withdrawn from the left ventricle and plasma separated. RBCs from these mice were washed thoroughly to remove free heme; RBC suspensions of 1.0% hematocrit along with 4mm solid glass beads (Pyrex) in DPBS were rotated 360° for 2 hours at 24 rpm at 37°C. The hemoglobin released from the RBCs during rotation was transferred into a new tube and centrifuged at 13,400g for 4 min and the absorbance of the supernatant were recorded at 540nm. Means ± SEM; n=8 for each group **(A)**. Next, total heme levels was measured in the plasma of air and COCl_2_ exposed mice using the QuantiChrom heme assay kit, according to the manufacturer’s instructions. Means ± 1 SEM; n=16 air and 20 for COCl_2_. (**B**). in the second series of experiments total (**C**) and free (**D**) heme levels in plasma were measured using an absorbance spectrum deconvolution (45). Means ± 1 SEM; n=33 air and 19 for COCl_2._ Total heme levels determined by these two methods were very similar.

### Exposure to COCl_2_ damages plasmalogens

Our previous data show that the halogens, Cl_2_ and Br_2_ interact with lung plasmalogens resulting in the formation of halogenated lipids (fatty acids and aldehydes), which cause extensive injury to RBCs and distant targets (10; 15; 46). Herein we show that exposure of mice to COCl_2_ (20 ppm for 10 min) results in significant decrease of plasmenylethanoamine **(Figure 9A)** and an increase of its breakdown product, polyunsaturated lysophosphatidylethanolamine **(Figure 9B)**.

**Figure 9.**
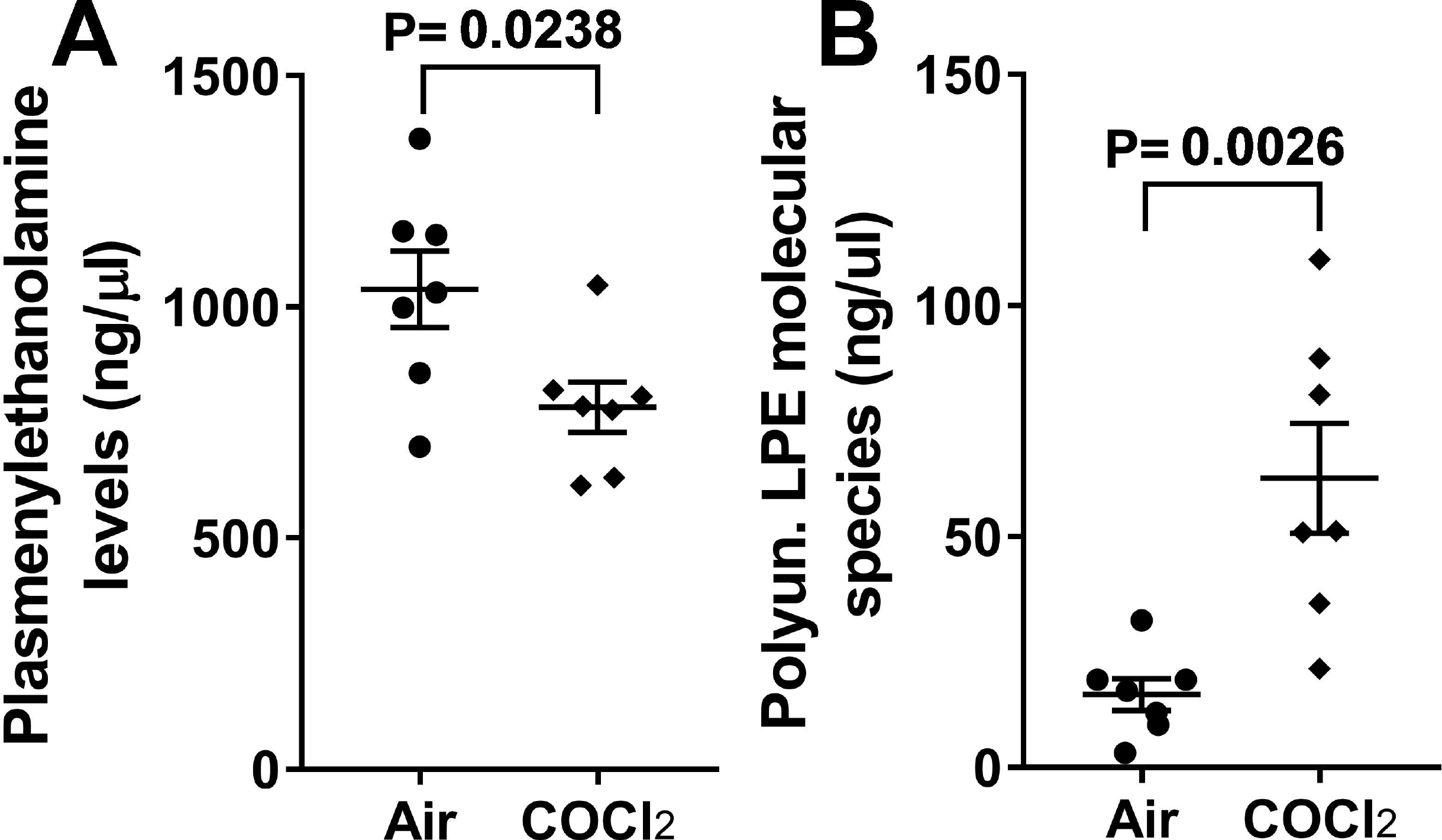
Measurement of plasmalogens and their breakdown product in COCl_2_ exposed mice. Equal number of male and female mice were sacrificed 24 h post exposure to either air or COCl_2_. Plasma was subjected to a modified Bligh-Dyer lipid extraction in the presence of lipid class internal standards including 1-0-tetradecanoyl-sn-glycero-3-phosphoethanolamine and 1,2-ditetradecanoyl-sn-glycero-3-phosphoethanolamine. Lipid extracts were diluted in methanol/chloroform (1/1, v/v) and molecular species were quantified using electrospray ionization mass spectrometry on a triple quadrupole instrument employing shotgun lipidomics methodologies (see Material and Methods for more details). Plasmenylethanolamine, **(A)** and Polyunsaturated Lysophosphatidylethanolamine **(B)** levels were measured. Values are means ± 1 SEM. Statistical Analysis by the Student’s t-test

## Discussion

Herein we demonstrate that male and female unanesthetized mice exposed to 20 pmm of COCl_2_ for 10 min and returned to room air, exhibited significant mortality with a LC_50_ of about 5 days. At 24 h post exposure there was significant increase of airway resistance and extensive injury to the blood gas barrier, resulting in the filling of large number of alveoli with fibrinous exudate, resulting in severe hypoxemia, hypercapnia and partially compensated respiratory acidosis. In addition, we observed significant decreases of plasma plasmalogens, with concomitant increase of plasmalogens degradation products, as well as oxidation of key structural and functional proteins of RBCs which may be responsible at least in part for increased RBC fragility. Furthermore, we documented the presence of free hemoglobin and heme in the plasma which may be responsible for initiation and propagation of injury to distal organs. Exposure to 10 ppm of COCl_2_ for 10 min did not result in mortality up to 21 days post exposure and the extent of lung injury was milder than observed with exposure to 20 ppm. There were no differences in any of the measured variables among male and female mice, although a large number of additional studies are needed to fully document this.

To this extent, the results of these studies mimic what has been found in humans. COCl_2_-induced ALI show evidence of persistent apnea periods, bradycardia, and higher pulmonary edema compared to Cl_2_ toxicity (35). At higher concentrations (>150 ppm·min) COCl_2_ exposure can cause life-threatening and latent non-cardiogenic pulmonary edema that is seen 6 to 24 hours post-exposure (11; 47). Patients who survive ALI from any source often develop chronic lung disease, with airflow obstruction, fibrosis (43), airway hyper-reactivity, and impaired gas exchange (5; 25; 50). These patients often require hospitalization (41; 52), and are predisposed to bacterial infections(8). A severe accident involving release of COCl_2_ occurred in a DuPont Factory in Belle, West Virginia in 2010 resulting in worker fatalities. Phosgene may also be generated during fires involving plastics and solvents containing Cl_2_ placing workers and first responders at serious risk (53). According to the Medical Management Guidelines for Phosgene issued by the Agency for Toxic Substances Disease Registry, there is currently no effective medical countermeasure for phosgene exposure, and emergency medical treatment (like in the case of Cl_2_ and Br_2_) consists of support of cardiopulmonary functions via supplemental oxygen and possibly bronchodilators and corticosteroids https://www.atsdr.cdc.gov/mmg/mmg.asp?id=1201&tid=182

In this study, we opted to expose mice to COCl_2_ in a whole body manner, instead of nose only, for various reasons: first we believe that whole body exposure better mimics human exposures; second, we have previously shown that exposure of mice to Cl_2_ gas causes the upregulation of the unfolded protein response which may result in lung injury, compounding the effects of the toxic gas (34); third, mice exposed in a nose only way are restrained in contrast to our system where they can move around freely. We carefully measured COCl_2_ concentrations in exposure chambers allowing the accurate calculation of the *ppm x time* profile which may be of considerable use to compare results of studies between various laboratories.

The mechanisms responsible for COCl_2_ induced injury are poorly understood. COCl_2_ is a very reactive gas and like Cl_2_ and Br_2_ increase the concentration of reactive species in the cytoplasm and mitochondria (4; 13; 31). Recently, we have demonstrated that Cl_2_ and Br_2_ react with plasmalogens, present in all cell membranes and the epithelial lining fluid, as well as glutathione to form halogenated adducts (fatty acids and aldehydes), which were detected in the alveolar space, lung tissues, plasma and red blood cell membranes (10; 15; 29). These are potentially damaging species which may destabilize cell membranes and cause extensive injury to the blood gas barrier. Phosgene also reacts with phosphatidylcholine, an important component of cell lipid membranes and pulmonary surfactant (18). Decreased levels of surfactant lipids as well as damage to surfactant proteins may result in severe lung injury, characterized by permeability-type edema and hypoxemia (20; 24). In the present study we found that plasma levels of plasmenylethanolamine, one of the most common plasmalogens, were decreased and plasma levels of lysophosphatidylethanolamine, a partial hydrolysis product of plasmenylethanolamine were increased in COCl_2_ exposed mice as compared to air exposed mice, suggesting that COCl_2_ may cross the blood-gas barrier and react with plasma plasmalogens. In addition, HCl, a COCL_2_ byproduct, has been shown to cause acute lung injury when instilled into the lungs of mice (14; 59). Injury may be compounded by the resulting inflammation due to the egress of neutrophils into the alveolar space. In this respect, in the present study we found that plasma levels of plasmenylethanolamine, one of the most common plasmalogens, were decreased and plasma levels of lysophosphatidylethanolamine, a partial hydrolysis product of plasmenylethanolamine were increased in COCl_2_ exposed mice as compared to air exposed mice, suggesting that COCl_2_ reacts with plasmalogens as well.

Exposure of mice to Cl_2_ or Br_2_ at 600 ppm for 30 minutes cause approximately 50% mortality within 7 days, similarly to 20 ppm COCl_2_ for 10 minutes (20/10) in the present study. We typically found BAL protein levels of 1 mg/mL to 1.5 mg/mL in Cl_2_ and Br_2_ exposed mice (1; 2; 26–28; 32; 57; 58), as compared to the typical 0.3 to 0.5 mg/mL protein levels in air exposed mice. Strikingly, we observed 10.3 mg/mL mean protein levels in BAL of mice exposed to COCl_2_, as compared to the 0.5 mg/mL mean protein levels in air exposed mice. Another striking difference is that most alveoli of COCl_2_ mice at 24 h post exposure were filled with fibrinous exudate; in contrast Cl_2_ and Br_2_ induced injury is patchier. Similarly, mean lung wet/dry ratios in mice exposed to Br_2_ and Cl_2_ range from 4.5 to 4.8, as compared to ~4.2-4.3 (ibid) in air exposed mice, but we found mean lung wet/dry ratios of 5.1, with individual values being as high as 6.5 in COCl_2_ exposed mice. These findings suggest that exposure conditions that lead to similar mortality outcomes are associated a significantly greater lung injury in COCl_2_ exposed mice, compared to Cl_2_ or Br_2_. The reasons for this difference are yet to be determined.

Arterial blood gas measurements in mice are relatively uncommon, due to the difficulty of collecting pure arterial blood in sufficient quantities. P_a_O_2_ and P_a_CO_2_ values in air-exposed mice, obtained while they were anesthetized, were higher and lower respectively compared to what has been reported in conscious mice (33). Following COCl_2_ exposure, P_a_O_2_ levels were decreased and P_a_CO_2_ levels were increased. There was a surprisingly adequate metabolic compensation with increase in 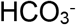 concentrations which returned the pH towards a normal value. These data indicate that the observed metrics of lung injury correlate well with the observed deficiency in gas exchange.

In addition to direct injury to the lung, halogens cause indirect injury to the lungs and systemic organs by causing red blood cell hemolysis (1; 2). Both heme and hemoglobin (Hb), released post hemolysis, are potent pro-oxidants and pro-inflammatory mediators (16; 23). Heme-induced lipid peroxidation (23) and oxidative stress damages the sugar-phosphate backbone causing strand breaks in the mitochondrial DNA (49). Elevated free hemoglobin/heme levels are associated with end organ injury and mortality in critically ill patients and therapies that free heme and hemoglobin are protective in animal models where hemolysis occurs(1; 2; 30; 36; 38; 39; 54)

In summary, these findings characterize in detail the development of lung and systemic injury in mice exposed to COCl_2_, offer significant new insights into the mechanisms leading to RBC fracture and release of free hemoglobin and heme and set the stage for the development of new countermeasures to limit lung injury and prolong survival.

## Authors Contributions

**SA, TJ**: Study design, data acquisition, analysis, interpretation, and writing of the manuscript; **SD, IA, SG, SE,CJA, J-Y O**: Data acquisition and analysis, and interpretation; **RPP, MG, DF:** Study design, analysis, data interpretation, and quality control. **SM**: Study design, analysis, interpretation, writing of the article, and responsible for quality control.

## Acknowledgements

**Funding**: Supported by the CounterACT Program, National Institutes of Health Office of the Director (NIH OD), the National Institute of Neurological Disorders and Stroke (NINDS), and the National Institute of Environmental Health Sciences (NIEHS), Grant Numbers (5UO1 ES026458 03; 3UO1 ES026458 03S1; 5UO1 ES027697 02) to SM. The authors would like to thanks Drs. Wesley W. Holmes, Alfred M. Sciuto and Dana R. Anderson from the Analytical Toxicology Division, USA Army Medical Research Institute of Chemical Defense for many useful discussions on phosgene toxicology.

